# AlphaFold 3-powered discovery of phage proteins that inhibit bacterial transcription

**DOI:** 10.64898/2025.12.17.695035

**Authors:** Linggang Yuan, Qingyang Liu, Xiaojian Xiao, Liqiao Xu, Liang Liang, Yang Guo, Yue Yao, Youjun Feng, Xiaoting Hua, Yu Feng

**Affiliations:** Department of Infectious Diseases, Sir Run Run Shaw Hospital, Zhejiang University School of Medicine, Hangzhou, China; Department of Biophysics, Zhejiang University School of Medicine, Hangzhou, China; Department of General Intensive Care Unit of the Second Affiliated Hospital, Zhejiang University School of Medicine, Hangzhou, China; Key Laboratory for Diagnosis and Treatment of Physic-Chemical and Aging Injury Diseases of Zhejiang Province, Hangzhou, China; State Key Laboratory of Transvascular Implantation Devices, Hangzhou, China

**Keywords:** AlphaFold 3, transcription regulation, phage, RNA polymerase, σ factor

## Abstract

Phages are the most abundant biological entities on Earth and play central roles in bacterial evolution and the emergence of new pathogens. Many phages encode proteins that specifically inhibit host RNA polymerase activity, thereby sabotaging and, in some cases, hijacking the host transcription machinery to serve their needs. Identification and characterization of these transcription inhibitors not only provide insights into the logic of transcription regulation but also inspire the design of antibiotics targeting bacterial transcription. Traditional methods for identifying new phage proteins that inhibit bacterial transcription are labor-intensive and require access to live phages. To overcome these limitations, we developed a highly efficient pipeline for AlphaFold 3–guided discovery of phage proteins that inhibit bacterial transcription. Using this pipeline, three phage proteins were identified and characterized. Structural and biochemical analyses demonstrated that these phage proteins bind to distinct sites on RNA polymerase and inhibit transcription *via* unprecedented mechanisms. This study showcases the power of AlphaFold 3 in discovering novel binders of large protein complexes, and the pipeline developed here could be readily adapted to screen modulators of other large targets, such as the ribosome, proteasome, and CRISPR-Cas systems.

## Introduction

Transcription is catalyzed by RNA polymerase (RNAP), a multi-subunit protein machinery. All bacterial RNAPs contain two copies of the α subunit and one copy each of the β, β′, and ω subunits ^1^. In addition to these conserved subunits, RNAPs from *Firmicutes* possess two additional subunits, δ and ε ^2–9^. The overall shape of RNAP resembles a crab claw with two pincers ^10,11^. These pincers form the main channel that accommodates double-stranded DNA (dsDNA), and the rotation of one pincer (the clamp) leads to the opening and closing of this channel ^12,13^. The catalytic center is located deep inside the crab-claw structure. Although the bacterial RNAP core enzyme can elongate RNA by adding nucleotides to the 3′ end of the nascent transcript, it cannot initiate transcription from promoter DNA on its own. To recognize promoters and initiate *de novo* transcription, the core enzyme must associate with an σ factor to form the RNAP holoenzyme ^14–16^. Bacterial genomes encode multiple σ factors, among which σ^A^ (σ^70^ in *E. coli*) is responsible for the transcription of most genes. σ^A^ typically consists of several conserved domains connected by flexible linkers ^17^. In the absence of promoter DNA, the σ_1.1_ domain occupies the main channel, thereby preventing nonspecific DNA binding ^18^. Upon promoter binding, σ_1.1_ is displaced, vacating the channel for promoter DNA entry. Within the holoenzyme, σ_2_ and σ_3_ interact with the clamp and recognize the −10 and extended −10 promoter elements, respectively, while σ_4_ is positioned at the RNA exit channel and binds the −35 element ^19–27^. The linker region (σ_3.2_) connecting σ_3_ and σ_4_ extends into the catalytic center, guiding the template-strand DNA within the transcription bubble and pre-organizing it into an A-form conformation.

Many phages encode transcription factors that specifically inhibit host RNAP, thereby sabotaging or even co-opting the transcription machinery to serve viral needs ^28–30^. These factors share neither sequence nor structural similarity, bind to distinct sites on RNAP, and employ diverse inhibitory strategies. For instance, the Ocr protein of phage T7 competes with σ factors for binding to the RNAP core enzyme ^31^, while its Gp2 protein anchors itself in the main channel by contacting both the RNAP jaw and σ_1.1_, thereby blocking promoter DNA entry ^32–35^. Likewise, the Gp39 protein of *Thermus thermophilus* phage P23-45 displaces σ_4_ to disrupt recognition of the −35 promoter element, while its Gp76 protein targets σ_3.2_ to preclude the engagement of the template-strand DNA of the transcription bubble ^36,37^. Similarly, the AsiA protein of T4 phage remodels σ_4_ to interfere with −35 element recognition ^38–41^. Given the vast diversity of phage protein space, it is likely that many more, yet-undiscovered phage proteins inhibit bacterial transcription through entirely novel mechanisms.

In this work, we establish a computational–experimental pipeline that leverages AlphaFold 3 to discover previously uncharacterized phage proteins that inhibit bacterial RNAP. Using this approach, we identified three phage proteins (EP1, PP1, and VP1) with potent inhibitory activity. Cryo-electron microscopy (Cryo-EM) structural analyses revealed that EP1 and PP1 displace σ_4_ by binding outside the RNA exit channel, while VP1 bridges the β subunit and σ factor to block open complex formation. These findings not only expand the catalog of phage transcription inhibitors but also demonstrate the power of artificial intelligence–guided discovery of protein modulators for large macromolecular machines.

## Results

### AlphaFold 3–guided discovery of phage inhibitor of bacterial RNAP

After rounds of optimization, we established a highly efficient pipeline for AlphaFold 3–guided discovery of phage proteins that inhibit bacterial transcription (Fig. 1a). To reduce computational demand, we leveraged UniProt Reference Clusters (UniRef), which provide non-redundant sets of sequences from UniProtKB and selected UniProt Archive records ^42^. Specifically, UniRef50, which groups sequences with ≥50% identity and ≥80% overlap with the longest sequence in the cluster, was used as the screening pool. Only clusters corresponding to phage proteins of unknown function were included. Because most phage transcription inhibitors identified to date are shorter than 200 amino acids, clusters exceeding this length were excluded. To avoid sequencing artifacts, clusters with fewer than 10 sequences were also removed. In total, 1,280 clusters of *Escherichia* phage proteins (Supplementary Table 1), 887 clusters of *Pseudomonas* phage proteins (Supplementary Table 2), and 512 clusters of *Vibrio* phage proteins (Supplementary Table 3) were retained. For each cluster, a representative protein was modeled in complex with the RNAP holoenzyme using AlphaFold 3 ^43^. Because RNAP is much larger than the phage proteins, the overall scores of the complexes were consistently ∼0.8. However, the interface predicted template modeling (ipTM) scores for the phage protein chain varied substantially (Fig. 1b). Since ipTM reflects the accuracy of the predicted relative positioning of the phage protein within the complex, we found it to be a strong predictor of true binding. Clusters were therefore selected for experimental validation only when the ipTM score exceeded 0.4 and the five predicted complex models were consistent. Based on these criteria, five *Escherichia* phage genes, three *Pseudomonas* phage genes, and one *Vibrio* phage gene were synthesized and cloned into recombinant expression vectors. Of these, four *Escherichia* phage proteins, two *Pseudomonas* phage proteins, and one *Vibrio* phage protein were successfully expressed in *E. coli* (BL21) and purified by affinity chromatography.

**Fig. 1:**
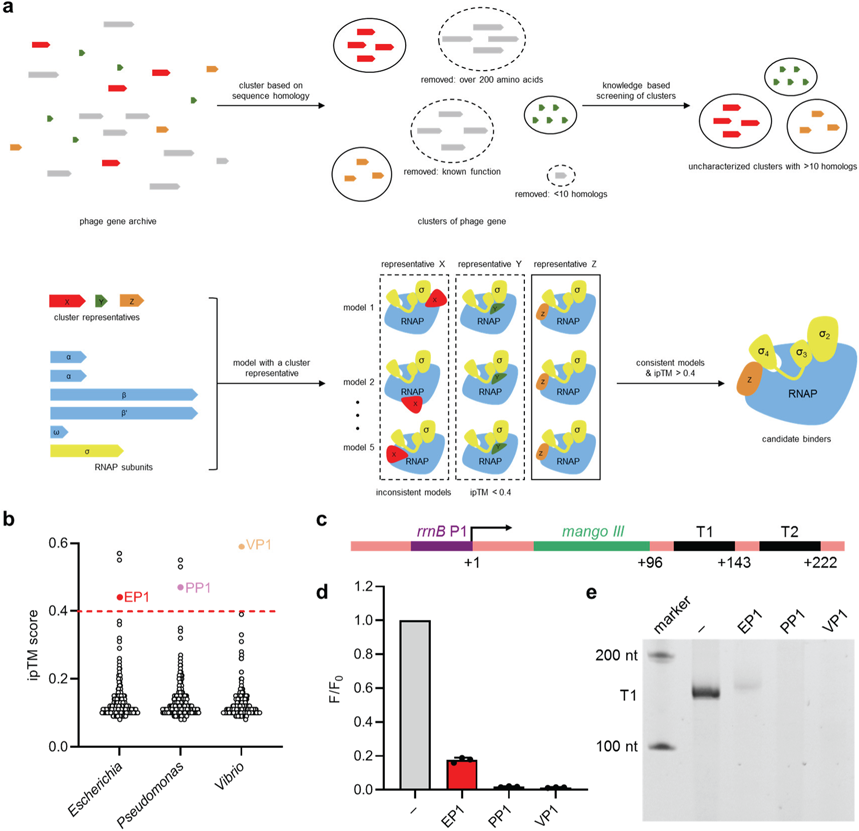
AlphaFold 3-guided discovery of phage inhibitor of bacterial RNAP. **a,** Pipeline for AlphaFold 3-guided discovery of phage inhibitor of bacterial RNAP. **b,** Distribution of interface predicted template modeling (ipTM) scores. The dashed line represents the 0.4 cutoff. Highlighted are phage proteins that demonstrated transcription inhibition activity. **c,** The DNA template for *in vitro* transcription assays. The DNA template contains *rrnB* P1 promoter followed by the Mango III sequence and *rrnB* terminators (T1 and T2). The transcription start site is highlighted by an arrow. **d,** Fluorescent transcription assay confirmed the transcription inhibition activity. 20 nM RNAP holoenzyme was incubated with 1 μM phage proteins and 20 nM promoter DNA for 10 min at 37°C before the addition of NTPs and TO1-Biotin. F, fluorescence intensity with phage proteins; F_0_, fluorescence intensity without phage proteins. **e,** Urea-PAGE transcription assay verified the transcription inhibition activity. 0.25 μM RNAP holoenzyme was incubated with 2 μM phage proteins and 0.1 μM DNA template for 10 min at 37°C before the addition of NTPs. The major products are approximately 150 nucleotides in length, consistent with the position of the *rrnB* T1 terminator.

To test their transcription inhibitory activity, we performed an *in vitro* transcription assay using the fluorogenic RNA aptamer Mango III ^44^. The DNA templates contained the *E. coli rrnB* P1 promoter, the Mango III sequence, and *rrnB* terminators (Fig. 1c). RNAP activity in the presence of each phage protein was quantified by fluorescence intensity of Mango III. Using this approach, one *Escherichia* phage protein (EP1, UniProtKB: A0A0N9RSC1), one *Pseudomonas* phage protein (PP1, UniProtKB: A0A0A1IUJ9), and one *Vibrio* phage protein (VP1, UniProtKB: A0A9X9HRU3) were confirmed to inhibit transcription by more than 50% at 1 μM (Fig. 1d). The inhibitory activities of these proteins were further validated by urea-PAGE analysis of transcription products (Fig. 1e). In both fluorescence- and gel-based assays, PP1 and VP1 exhibited stronger inhibition than EP1.

### Mechanism of transcription inhibition by EP1

To elucidate the structural basis of transcription inhibition by EP1, we incubated *E. coli* RNAP holoenzyme with a twofold molar excess of EP1 and determined the RNAP–EP1 complex structure by cryo-EM (Supplementary Fig. 1 and Table 4). The final reconstruction yielded a 4.0 Å map, sufficient to resolve protein secondary structures. Clear densities were observed for EP1 and the RNAP core enzyme, but no density corresponding to σ^70^ was detected (Fig. 2a and b). Compared with the RNAP holoenzyme (PDB: 6C9Y), the RNAP clamp in the EP1 complex is rotated by ∼11°, although less open than in the RNAP core enzyme (PDB: 7MKP, ∼23°, Fig. 2c). EP1 binds outside the RNA exit channel, contacting the N-terminal helix and zinc-binding domain of the β′ subunit and the C-terminus of the ω subunit, burying a total surface area of 1,403 Å^2^.

**Fig. 2:**
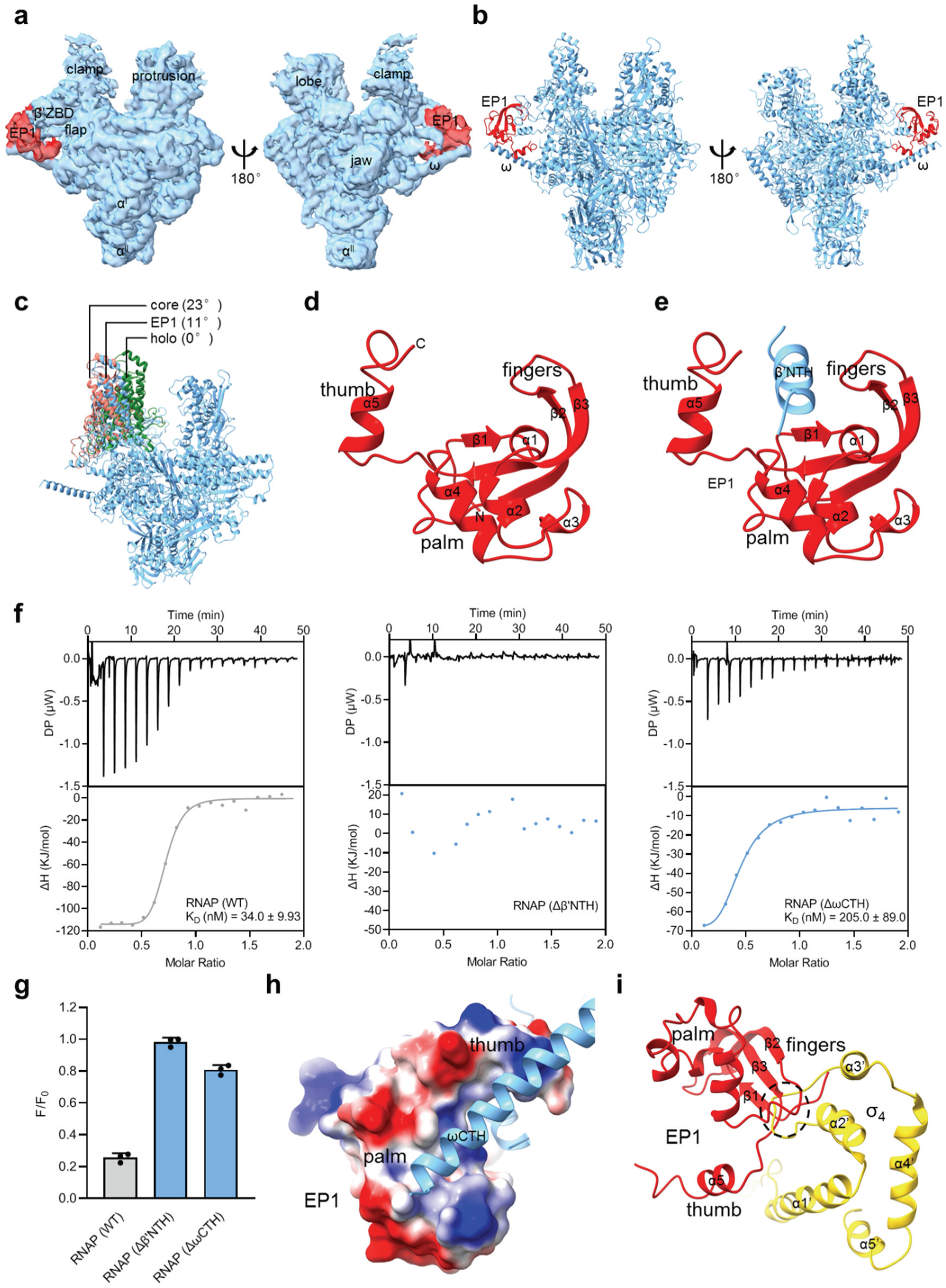
Mechanism of transcription inhibition by EP1. **a,** The overall map of RNAP-EP1. The map without B-factor sharpening is shown in opposite orientations. Cyan, RNAP core enzyme; red, EP1. **b,** The overall structure of RNAP-EP1. The structure is shown as ribbons. The view orientations and color scheme are the same as Fig. 2a. **c,** The RNAP clamp in the presence of EP1 is open relative to that of RNAP holoenzyme, but not as open as RNAP core enzyme. The structures are shown as ribbons. Cyan, RNAP-EP1; green, RNAP holoenzyme (PDB: 6C9Y); salmon, RNAP core enzyme (PDB: 7MKP). **d,** The structure of EP1 resembles a left hand. α1–α4 form the palm, while α5 and β1–β3 constitute the thumb and fingers, respectively. **e,** EP1 holds the N-terminal helix of β′ subunit with its palm, thumb, and fingers. β′NTH, β′ N-terminal helix. **f,** Isothermal titration calorimetry (ITC) experiments of RNAP derivatives. EP1 was gradually titrated to RNAP or its derivatives, and the heat released upon complex formation was precisely measured. β′NTH, β′ N-terminal helix; ωCTH, ω C-terminal helix. **g,** Fluorescent transcription assay of RNAP derivatives. 20 nM RNAP holoenzyme or its derivatives was incubated with 80 nM EP1 and 20 nM promoter DNA for 10 min at 37°C before the addition of NTPs and TO1-Biotin. F, fluorescence intensity with phage proteins; F_0_, fluorescence intensity without EP1. β′NTH, β′ N-terminal helix; ωCTH, ω C-terminal helix. **h,** The C-terminus of ω subunit adopts a helical conformation and binds to a positively charged groove in the palm. ωCTH, ω C-terminal helix. **i,** EP1 clashes with σ_4_. The structure of RNAP-EP1 is superimposed on the cryo-EM structure of *E. coli* RNAP holoenzyme (PDB: 6C9Y) based on the clamp residues.

EP1 adopts a unique fold composed of five α-helices (α1–α5) and three antiparallel β-strands (β1–β3) (Fig. 2d and Supplementary Fig. 2a). The overall structure resembles a left hand: α1–α4 form the palm, while α5 and β1–β3 constitute the thumb and fingers, respectively. A Dali search ^45^ revealed no structural homologs, indicating a previously undescribed protein fold. The “left hand” clamps the N-terminal helix of the β′ subunit with its palm, thumb, and fingers (Fig. 2e). Isothermal titration calorimetry (ITC) experiments demonstrated that EP1 binds RNA polymerase with a dissociation constant (K_D_) of 34 nM (Fig. 2f). Truncation of the β′ N-terminal helix (β′NTH) abolished the RNAP–EP1 interaction, indicating that β′NTH is the primary anchoring site of EP1. Consistently, truncation of β′NTH also abolished EP1-mediated transcription inhibition (Fig. 2g). In addition, the C-terminus of the ω subunit—which is disordered in the RNAP holoenzyme structure—adopts a helical conformation that inserts into a positively charged groove in the EP1 palm (Fig. 2h). Deletion of the ω C-terminal helix (ωCTH) reduced both binding affinity and inhibitory activity of EP1, though to a lesser extent than β′NTH truncation, suggesting that the ωCTH–EP1 interaction is important but not essential (Fig. 2f and g). In the RNAP–EP1 structure predicted by AlphaFold 3, EP1 adopts the same left-hand fold and clamps β′NTH as observed in the cryo-EM structure (Supplementary Fig. 3a). However, AlphaFold 3 failed to predict RNAP clamp opening and the EP1–ωCTH interaction, underscoring the importance of experimental validation.

To explain the absence of σ^70^ density, we superimposed the RNAP–EP1 complex onto the RNAP holoenzyme structure (PDB: 6C9Y) based on clamp residues (Fig. 2i). This revealed severe steric clashes between the EP1 “fingers” (β1–β3) and σ^70^ region 4 (σ_4_). We therefore conclude that EP1 inhibits transcription by displacing σ_4_ from RNAP.

### Mechanism of transcription inhibition by PP1

To investigate the mechanism of transcription inhibition by PP1, we determined the cryo-EM structure of the *E. coli* RNAP–PP1 complex using the same approach as for RNAP–EP1 (Supplementary Fig. 4 and Table 4). Well-resolved densities were obtained for both PP1 and the RNAP holoenzyme, with the exception of σ^70^ (Fig. 3a and b). Compared with the RNAP holoenzyme, the clamp in the RNAP–PP1 complex is rotated by ∼17°, an even greater shift than that observed in the RNAP–EP1 complex (Fig. 3c). PP1 binds outside the RNA exit channel, contacting the N-terminal helix, zinc-binding domain, dock, lid, and zipper of the β′ subunit, as well as the C-terminal region of the β subunit, burying a surface area of 2,624 Å^2^—nearly twice that of EP1. Although AlphaFold 3 correctly predicted the folding of PP1 and its binding position, it again failed to predict the rotation of the RNAP clamp (Supplementary Fig. 3b).

**Fig. 3:**
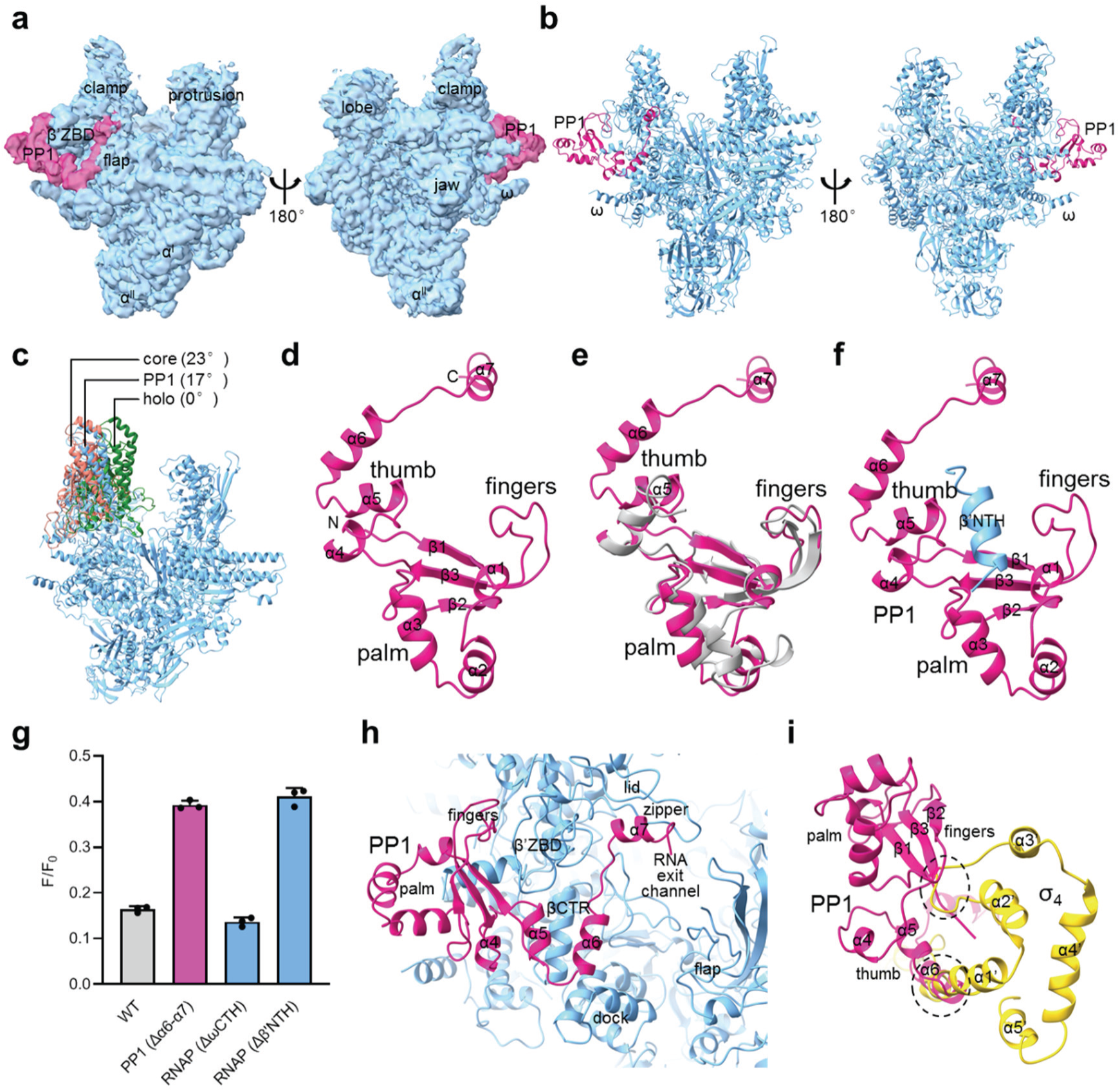
Mechanism of transcription inhibition by PP1. **a,** The overall map of RNAP-PP1. The map without B-factor sharpening is shown in opposite orientations. Cyan, RNAP core enzyme; magenta, PP1. **b,** The overall structure of RNAP-PP1. The structure is shown as ribbons. The view orientations and color scheme are the same as Fig. 3a. **c,** The clamp in the presence of PP1 is more open than that of RNAP-EP1. The structures are shown as ribbons. Cyan, RNAP-PP1; green, RNAP holoenzyme (PDB: 6C9Y); salmon, RNAP core enzyme (PDB: 7MKP). **d,** The structure of PP1 resembles a left hand, and the thumb is extended with three helices. α1–α3 form the palm, while α4–α7 and β1–β3 constitute the thumb and fingers, respectively. **e,** Compared with EP1, the palm of PP1 has fewer helices and the thumb of PP1 is extended with three extra helices. Magenta, PP1; gray, EP1. **f,** PP1 holds the N-terminal helix of β′ subunit with its palm, thumb, and fingers. β′NTH, β′ N-terminal helix. **g,** Fluorescent transcription assay of PP1 and RNAP derivatives. 20 nM RNAP holoenzyme was incubated with 40 nM PP1 or its derivative and 20 nM promoter DNA for 10 min at 37°C before the addition of NTPs and TO1-Biotin. F, fluorescence intensity with PP1 or its derivative; F_0_, fluorescence intensity without PP1. β′NTH, β′ N-terminal helix; ωCTH, ω C-terminal helix. **h,** The C-terminal helix of PP1 invades the RNA exit channel and contacts the lid and zipper of β′ subunit. β′ZBD, β′ zinc-binding domain; βCTR, β C-terminal region. **i,** PP1 clashes with σ_4_. The structure of RNAP-PP1 is superimposed on the cryo-EM structure of *E. coli* RNAP holoenzyme (PDB: 6C9Y) based on the clamp residues.

PP1 adopts a fold comprising seven α-helices (α1–α7) and three antiparallel β-strands (β1–β3, Fig. 3d, 3e, and Supplementary Fig. 2b). Its tertiary structure also resembles a left hand, with the palm (α1–α3), thumb (α4-α7), and fingers (β1–β3). As in EP1, this “hand” grips the N-terminal helix of the β′ subunit (Fig. 3f), and truncation of this helix significantly reduced PP1-mediated inhibition (Fig. 3g). However, unlike EP1, the palm of PP1 contains fewer helices and does not contact the ω subunit C-terminus; accordingly, ω truncation had no effect on inhibition (Fig. 3g). Distinctly, PP1 possesses an extended thumb formed by α4–α7. Notably, α5 and α6 fold back to position α7 deep within the RNA exit channel, where it engages the lid and zipper of the β′ subunit (Fig. 3h). Consistent with this structural role, truncation of α6–α7 markedly decreased inhibitory activity (Fig. 3g).

PP1 also inhibits transcription by displacing σ^70^ region 4 (σ_4_). Structural alignment revealed steric clashes between the PP1 “fingers” (β1–β3) and σ_4_, while α6 of PP1 occupies the same spatial position as the first helix of σ_4_ (Fig. 3i). Consequently, disrupting either the β′ N-terminal helix interaction or α6–σ_4_ overlap reduced—but did not abolish—PP1-mediated inhibition (Fig. 3g).

### Mechanism of transcription inhibition by VP1

To determine the structural basis of VP1-mediated transcription inhibition, we solved the cryo-EM structure of the *E. coli* RNAP holoenzyme in complex with VP1 at 3.3 Å resolution (Supplementary Fig. 5 and Table 4). Unlike in the EP1 and PP1 complexes, σ^70^ density was fully retained (Fig. 4a and b), and the clamp adopted a closed conformation similar to that of the RNAP holoenzyme (Fig. 4c). VP1 binds within the main channel, bridging the β-subunit protrusion and σ^70^ region 2 (σ_2_), burying a total surface area of 1,604 Å^2^. The local resolution of ∼3 Å enabled detailed analysis of the interacting residues. The structure predicted by AlphaFold 3 closely matches the cryo-EM structure, with a root-mean-square deviation (RMSD) of 0.788 Å (Supplementary Fig. 3c).

**Fig. 4:**
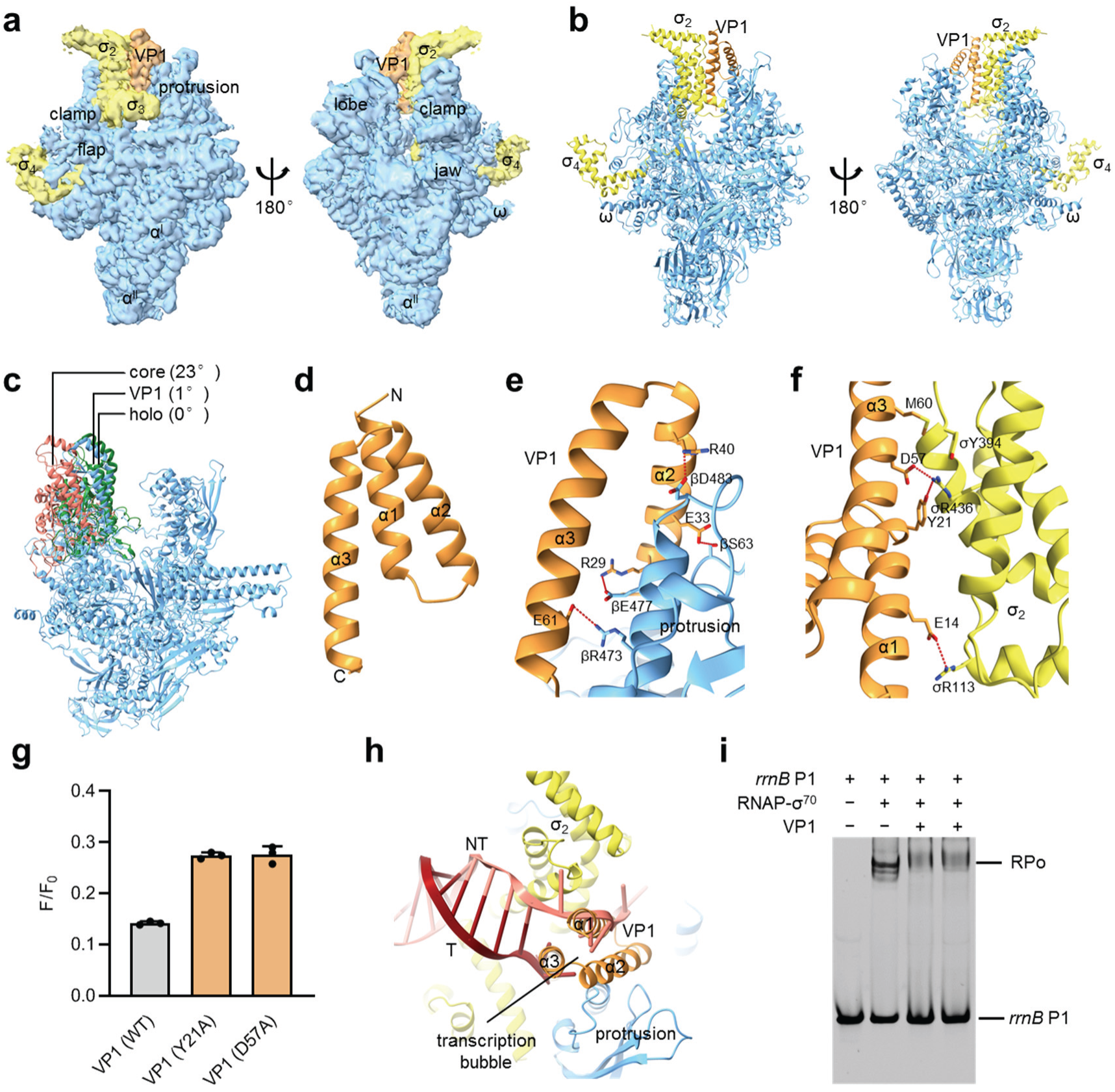
Mechanism of transcription inhibition by VP1. **a,** The overall map of RNAP-VP1. The map without B-factor sharpening is shown in opposite orientations. Cyan, RNAP core enzyme; yellow, σ^70^; orange, PP1. **b,** The overall structure of RNAP-VP1. The structure is shown as ribbons. The view orientations and color scheme are the same as Fig. 4a. **c,** The clamp in the presence of VP1 is closed. The structures are shown as ribbons. Cyan, RNAP-VP1; green, RNAP holoenzyme (PDB: 6C9Y); salmon, RNAP core enzyme (PDB: 7MKP). **d,** VP1 consists of three α-helices, folding into a helical bundle. **e,** α2 and α3 of VP1 contact the protrusion of β subunit. Contact residues are shown as sticks. Salt bridges and hydrogen bonds are shown as dashed lines. **f,** α1 and α3 of VP1 contact σ_2_. Contact residues are shown as sticks. Salt bridges and hydrogen bonds are shown as dashed lines. **g,** Fluorescent transcription assay of VP1 derivatives. 20 nM RNAP holoenzyme was incubated with 40 nM VP1 or its derivatives and 20 nM promoter DNA for 10 min at 37°C before the addition of NTPs and TO1-Biotin. F, fluorescence intensity with phage proteins; F_0_, fluorescence intensity without VP1. **h,** Structural alignment of RNAP-VP1 and RPo (PDB: 7KHB) shows that α1 and α3 of VP1 occupy the position of the nontemplate- and template-strand DNA of the transcription bubble. NT, nontemplate-strand DNA; T, template-strand DNA. **i,** Gel shift assay demonstrated that VP1 significantly decreases the amount of RPo formed with promoter *rrnB* P1. Lane 1, 1.2 μM promoter DNA; lane 2, 1 μM RNAP holoenzyme was incubated with 1.2 μM promoter DNA for 10 min at 37°C; lane 3, 1 μM RNAP holoenzyme was incubated with 2 μM VP1 for 10 min at 37°C before the addition of 1.2 μM promoter DNA; lane 4, 1 μM RNAP holoenzyme was incubated with 1.2 μM promoter DNA for 10 min at 37°C before the addition of 2 μM VP1.

VP1 consists of three α-helices (α1–α3) that fold into a helical bundle (Fig. 4d and Supplementary Fig. 2c). α2 and α3 interact with the β subunit protrusion mainly through electrostatic interactions (Fig. 4e). Specifically, R29 and R40 on α2 form salt bridges with β residues E477 and D483, respectively, while E61 on α3 forms a salt bridge with β residue R473. In addition, the carboxyl group of E33 on α2 forms a hydrogen bond with the hydroxyl group of β residue S63. Besides interacting with the protrusion, VP1 also contacts σ_2_ through α1 and α3 (Fig. 4f). In particular, E14 on α1 and D57 on α3 make electrostatic interactions with σ residues R113 and R436, respectively. Moreover, the hydroxyl group of Y21 on α1 forms a hydrogen bond with the guanidinium group of σ residue R436, while M60 on α3 makes a van der Waals interaction with σ residue Y394. Consistent with the structural analyses, all interacting residues are highly conserved among VP1 homologs (Supplementary Fig. 6). Supporting this, alanine substitutions of residues Y21 and D57 impaired VP1-mediated transcription inhibition (Fig. 4g). Most β subunit and σ factor residues are conserved between *Escherichia* and *Vibrio*, explaining the potent inhibitory activity against *E. coli* RNA polymerase (Supplementary Figs. 7 and 8).

Structural analyses reveal a dual mechanism of inhibition. First, promoter DNA loading requires clamp rotation during RNAP–promoter open complex (RPo) formation ^46,47^, but VP1 locks the clamp closed by bridging the β protrusion and σ_2_. Second, structural alignment of the RNAP–VP1 complex with RPo (PDB: 7KHB) shows that α1 and α3 of VP1 sterically occupy the positions of the nontemplate- and template-strand DNA within the transcription bubble (Fig. 4h), thereby preventing bubble formation. Consistent with this model, gel-shift assays showed that VP1 markedly reduces RPo formation at the *rrnB* P1 promoter (Fig. 4i). Moreover, addition of VP1 to pre-formed RPo efficiently disrupted the complex, confirming its inhibitory activity.

## Discussion

In summary, we identified three phage inhibitors of bacterial RNAP using the predictive power of AlphaFold 3 and characterized their mechanisms through structural and biochemical analyses. Although EP1 and PP1 share little sequence similarity, both contain a “left-hand” domain that binds outside the RNA exit channel and sterically clashes with σ_4_. Notably, PP1 features an extended thumb of three helices that folds back to insert into the RNA exit channel, enlarging the interaction interface (2,624 Å^2^ *vs.* 1,403 Å^2^ for EP1) and enhancing inhibitory efficiency (Fig. 1d and e). By contrast, VP1 adopts a three-helix bundle that bridges two RNAP elements and occupies the position of the transcription bubble, thereby locking the clamp closed and blocking open-complex formation.

Phages have long been recognized as key drivers of bacterial evolution, and the discovery of novel transcription inhibitors further highlights their role in shaping host regulatory strategies. The fact that EP1 and PP1, despite lacking sequence similarity, both adopt a “left hand” fold to target σ_4_ suggests a striking example of convergent evolution, where distinct phages independently arrived at a common solution to disrupt promoter recognition. Similarly, the compact helical bundle of VP1 illustrates yet another evolutionary innovation to interfere with RNAP function by locking the clamp and blocking bubble formation. These findings underscore the remarkable structural diversity of phage proteins and their ability to exploit different vulnerabilities of the same essential host machinery.

Bacterial RNAP is a validated target for antimicrobial drug discovery ^48,49^. The best-known RNAP inhibitors in clinical use are rifampin and its analogs, which have served as first-line treatments for tuberculosis and biofilm-associated infections for decades. These drugs bind the β subunit and block RNA synthesis beyond three nucleotides ^10,50^. Another RNAP inhibitor in clinical use is fidaxomicin, which selectively targets *Clostridioides difficile* while sparing the gut microbiome ^51,52^. Structural and single-molecule FRET studies have shown that fidaxomicin inhibits transcription initiation by jamming the clamp motion required for RPo formation ^53,54^. The identification and mechanistic characterization of phage proteins that inhibit bacterial transcription provide new conceptual frameworks for RNAP-targeted drug discovery. Small molecules or peptides could be designed to mimic the inhibitory strategies of phage proteins—for example, a peptide modeled on the C-terminal extension of PP1 could displace σ_4_ and suppress transcription.

To date, only a small number of phage proteins have been characterized as RNAP inhibitors, most of which originate from model phages such as T4 and T7. Traditional approaches for identifying new transcription inhibitors—such as biochemical or genetic assays—are labor intensive and require access to viable phages. To overcome these limitations, we leveraged artificial intelligence (AI)–based structure prediction, which can now model protein complexes with remarkable accuracy. Recently, an AlphaFold-guided strategy was successfully applied to discover viral inhibitors of host immunity ^55^. In that study, AlphaFold-Multimer was used to screen phage proteins for interactions with bacterial defense systems, leading to the discovery of seven classes of Thoeris and CBASS inhibitors. Similarly, AlphaFold-Multimer enabled the identification of binders to sperm fertilization factors from ∼1,400 testis-expressed membrane proteins, including Tmem81, which forms a heterotrimer with the essential fertilization factors Izumo1 and Spaca6 ^56^. Notably, the Thoeris, CBASS, and fertilization complexes involve relatively small proteins (<500 amino acids) ^57–59^, whereas bacterial RNAP holoenzyme is far more complex, comprising at least six subunits and >4,000 amino acids in total. While AI-powered structure prediction has proven effective for small complexes, its utility had not been tested on macromolecular assemblies of this scale. Our results demonstrate that this strategy can indeed resolve large and flexible protein complexes. Looking ahead, this platform could be adapted to systematically screen for endogenous or exogenous regulators of other major prokaryotic and eukaryotic machines, such as the ribosome, proteasome, and CRISPR-Cas systems.

## Methods

### AlphaFold 3-guided screening of RNAP inhibitor

Uncharacterized phage protein clusters were retrieved from the UniRef50 archive on March 5^th^, 2025. Only clusters with more than 10 sequences and less than 200 amino acids were retrieved. The sequence of the representative of each cluster was submitted together with the sequences of RNAP subunits on the AlphaFold Server (https://alphafoldserver.com/). 5 models were generated for each protein complex. The models were downloaded from the server and manually checked for consistency. The best ipTM value for the chain of phage protein among the 5 models were analyzed. Phage proteins with 5 consistent models and ipTM value larger than 0.4 were tested experimentally.

### Expression and purification of phage proteins

The genes encoding candidate phage proteins were synthesized and subcloned to pET21a by GENEWIZ, Inc. Candidate phage proteins were prepared from *E. coli* strain BL21(DE3) (Invitrogen, Inc.) transformed with expression plasmids under the control of the bacteriophage T7 gene 10 promoter. Single colonies of the resulting transformants were used to inoculate 1 l LB broth containing 100 μg/mL ampicillin, cultures were incubated at 37 °C with shaking until OD_600_ = 0.6, were induced by addition of IPTG to 1 mM, and were incubated overnight at 16°C. Then cells were harvested by centrifugation (5,000 x g; 10 min at 4°C), resuspended in 20 ml lysis buffer (20 mM Tris-HCl, pH 8.0, 0.2 M NaCl, 5% glycerol, 1 mM DTT) and lysed using a JN-02C cell disrupter (JNBIO, Inc.). After centrifugation (20,000 x g; 30 min at 4°C), the supernatant was loaded onto a column of Ni-NTA beads (Smart-Lifesciences, Inc.) equilibrated with lysis buffer. The column was washed with 30 ml lysis buffer containing 40 mM imidazole and eluted with 5 ml lysis buffer containing 0.3 M imidazole. Phage protein derivatives were constructed using Quikchange Site-Directed Mutagenesis Kits (Agilent, Inc.) and purified in the same way as the wildtype protein.

### Expression and purification of *E. coli* σ^70^

*E. coli* strain BL21(DE3) (Invitrogen, Inc.) was transformed with plasmid pGEMD ^60^. Single colonies of the resulting transformants were used to inoculate 1 l LB broth containing 100 μg/mL ampicillin, cultures were incubated at 37°C with shaking until OD_600_ = 0.6, were induced by addition of IPTG to 1 mM, and were incubated 3 h at 37°C. Then cells were harvested by centrifugation (5,000 x g; 15 min at 4°C), resuspended in 20 ml lysis buffer (40 mM Tris-HCl, pH 7.9, 0.3 M KCl, 10 mM EDTA, 1 mM DTT, and 0.2% sodium deoxycholate) and lysed using a JN-02C cell disrupter (JNBIO, Inc.). After centrifugation (20,000 x g; 30 min at 4°C), the pellet was washed with 25 ml lysis buffer twice, resuspended in 6 M GuaHCl, and dialyzed against 1 l dialysis buffer (20 mM Tris-HCl, pH7.9, 0.2 M NaCl, 1 mM EDTA, and 5 mM β-mercaptoethanol) twice. After centrifugation (20,000 x g; 30 min at 4°C), the supernatant was loaded onto a HiTrap Q HP column (GE Healthcare, Inc.) equilibrated in Q buffer (10 mM Tris-HCl, pH7.9, 1 mM EDTA, 1 mM DTT, and 5% glycerol) and eluted with a 100 ml linear gradient of 0.3–0.5 M NaCl in Q buffer. Fractions containing *E. coli* σ^70^ were pooled and stored at –80°C. Yield was ∼50 mg/l, and purity was >95%.

### Expression and purification of *E. coli* RNAP core enzyme

*E. coli* RNAP core enzyme was prepared from *E. coli* strain BL21(DE3) (Invitrogen, Inc.) transformed with plasmid pIA900 ^61^. Single colonies of the resulting transformants were used to inoculate 5 l LB broth containing 100 μg/mL ampicillin, cultures were incubated at 37°C with shaking until OD_600_ = 0.6, were induced by addition of IPTG to 1 mM, and were incubated 3 h at 37°C. Then cells were harvested by centrifugation (5,000 x g; 15 min at 4°C), resuspended in 120 ml lysis buffer (50 mM Tris-HCl, pH 7.9, 0.2 M NaCl, 2 mM EDTA, 5% glycerol, and 5 mM DTT) and lysed using a JN-02C cell disrupter (JNBIO, Inc.). After poly(ethyleneimine) precipitation and ammonium sulfate precipitation, the pellet was resuspended in Ni-NTA buffer (10 mM Tris-HCl, pH 7.9, 0.5 M NaCl, and 5% glycerol) and loaded onto a column of Ni-NTA beads (Smart-Lifesciences, Inc.) equilibrated with the same buffer. The column was washed with 50 ml Ni-NTA buffer containing 20 mM imidazole and eluted with 25 ml Ni-NTA buffer containing 0.15 M imidazole. The eluate was diluted in Q buffer and loaded onto a HiTrap Q HP column (GE Healthcare, Inc.) equilibrated in Q buffer and eluted with a 100 ml linear gradient of 0.3-0.5 M NaCl in Q buffer. Fractions containing *E. coli* RNAP core enzyme were pooled and stored at –80°C. Yield was ∼2.5 mg/l, and purity was >95%. RNAP derivatives were constructed using Quikchange Site-Directed Mutagenesis Kits (Agilent, Inc.) and purified in the same way as the wildtype protein.

### Purification of *E. coli* RNAP-σ^70^ holoenzyme

*E. coli* RNAP core enzyme and *E. coli* σ^70^ were incubated in a 1:4 ratio for 1 h at 4°C. The reaction mixture was applied to a HiLoad 16/600 Superdex 200 column (GE Healthcare, Inc.) equilibrated in 10 mM HEPES, pH 7.5, and 50 mM KCl, and the column was eluted with 120 ml of the same buffer. Fractions containing *E. coli* RNAP-σ^70^ holoenzyme were pooled and stored at –80°C.

### Fluorescent transcription assay

A DNA fragment containing *rrnB* P1 promoter followed by Mango III coding sequence and *rrnB* terminators was synthesized and inserted into pUC57 (GENEWIZ, Inc.) ^62^. The DNA fragment were amplified by PCR, was purified using the FastPure Gel DNA Extraction Mini Kit (Vazyme, Inc.), and was stored at –20°C. Fluorescent transcription assay was performed in a 96-well microplate format. Reaction mixtures contained (100 μl): 20 nM RNAP holoenzyme, 20 nM DNA fragment, various concentrations of phage protein, 50 mM Tris-HCl, pH 8.0, 0.1 M KCl, 10 mM MgCl_2_, 1 mM DTT, and 5% glycerol. After incubation for 10 min at 37°C, 1 μM TO1-Biotin, 0.2 mM ATP, 0.2 mM UTP, 0.2 mM GTP, and 0.2 mM CTP were added to start transcription. Following incubation for 1 h at 37°C, fluorescence emission intensities were measured using a Biotek Synergy H1 microplate reader (Agilent, Inc.; excitation wavelength = 510 nm; emission wavelength = 535 nm).

### Urea-PAGE transcription assay

The urea-PAGE transcription assay was performed using the same DNA template as the fluorescent transcription assay. Reaction mixtures contained (20 μl): 0.25 μM RNAP holoenzyme, 0.1 μM DNA fragment, 2 μM phage protein, 50 mM Tris-HCl, pH 8.0, 0.1 M KCl, 10 mM MgCl_2_, 1 mM DTT, and 5% glycerol. After incubation for 10 min at 37°C, 1 mM ATP, 1 mM UTP, 1 mM GTP, and 1 mM CTP were added to start transcription. Following incubation for 1 h at 37°C, 1 μl DNaseI (Ambion, Inc.) and 1 μl 5 mM CaCl_2_ was added to digest the DNA templates. The transcription products were mixed with 20 μl loading buffer (8M urea and 0.1% bromophenol blue), boiled for 5 min at 95°C, applied to 10% urea-polyacrylamide slab gels (19:1 acrylamide/bisacrylamide), electrophoresed in 0.5 x TBE buffer (44.5 mM Tris-borate, pH 8.0, and 1 mM EDTA), and stained with 4S Red Plus Nucleic Acid Stain (Sangon Biotech, Inc.).

### Electrophoretic mobility shift assay

Template strand DNA and non-template strand DNA (Genscript, Inc.) were annealed at a 1:1 ratio in 20 mM Tris-HCl, pH 8.0, 0.2 M NaCl, 1 mM DTT, and 5% glycerol. To test whether VP1 inhibits RPo formation, 1 μM *E. coli* RNAP-σ^70^ holoenzyme was incubated with 2 μM VP1 for 10 min at 37°C. Then 1.2 μM promoter DNA was added and incubated for 10 min at 37°C. To test whether VP1 disrupts pre-formed RPo, 1 μM *E. coli* RNAP-σ^70^ holoenzyme was incubated with 1.2 μM promoter DNA for 10 min at 37°C. Then 2 μM VP1 was added and incubated for 10 min at 37°C. The reaction mixtures were applied to 5% polyacrylamide slab gels (29:1 acrylamide/bisacrylamide), electrophoresed in 0.5 x TBE buffer (44.5 mM Tris-borate, pH 8.0, and 1 mM EDTA), and stained with 4S Red Plus Nucleic Acid Stain (Sangon Biotech, Inc.).

### Cryo-EM grid preparation

To determine the structure of phage protein in complex with RNAP, *E. coli* RNAP-σ^70^ holoenzyme was incubated with a twofold molar excess of phage protein for 20 min at 37°C. Immediately before freezing, 8 mM CHAPSO was added to the sample. Quantifoil grids (R 1.2/1.3, Cu, 300) were glow-discharged for 120 s at 25 mA prior to the application of 3 μl of the complex, then plunge-frozen in liquid ethane using a Vitrobot (FEI, Inc.) with 95% chamber humidity at 4°C.

### Cryo-EM data acquisition and processing

The grids were imaged using a 300 kV Titan Krios equipped with a Falcon 4 direct electron detector (FEI, Inc.). Images were recorded with EPU in counting mode with a physical pixel size of 1.19 Å and a defocus range of 1.0-2.0 μm. Images were recorded with a 5.65 s exposure to give a total dose of 50 e/Å^2^. Subframes were aligned and summed using UCSF MotionCor2^63^. The contrast transfer function was estimated for each summed image using CTFFIND4 ^64^.

From the summed images, approximately 1,000 particles were manually picked and subjected to 2D classification in RELION ^65^. 2D averages of the best classes were used as templates for auto-picking in RELION. Auto-picked particles were manually inspected, then subjected to 2D classification in RELION. Poorly populated classes were removed. The remaining particles were 3D classified in RELION using a map of *E. coli* RNAP low-pass filtered to 40 Å resolution as a reference. 3D classification resulted in 4 classes, among which only one class has a clear density for RNAP and phage protein. CTF refinement and particle polishing were performed before final 3D refinement and postprocessing.

### Cryo-EM model building and refinement

The models of phage proteins in complex with RNAP predicted by AlphaFold 3 ^43^ were fitted into the cryo-EM density map using Chimera ^66^ and were adjusted in Coot ^67^. The coordinates were real-space refined with secondary structure restraints in PHENIX ^68^.

### Isothermal Titration Calorimetry (ITC) experiment

To minimize the difference of solvent, the buffer of EP1 and RNAP was changed to phosphate buffered saline (PBS) using the HiPrep 26/10 Desalting column (GE Healthcare, Inc.). ITC experiments were performed with Microcal PEAQ-ITC (Malvern, Inc.) at 25°C. The cell is filled with 280 μl of 5 μM RNAP or its derivatives, and the syringe is filled with 38 μl of 50 μM EP1. A total of 19 titrations of EP1 with an interval time of 150 s were carried out. The first titration of 0.4 μl EP1 was carried out for 0.8 s. Then the other titrations of 2 μl EP1 were carried out for 4 s. The dissociation constant (K_D_) was fitted using PEAQ-ITC Analysis software v1.52 (Malvern, Inc.).

### Statistics and reproducibility

Statistics were performed in GraphPad Prism 8.0.2. No statistical method was used to predetermine sample size. No data were excluded from the analyses. The experiments were not randomized.

### Data availability

The cryo-EM density maps generated in this study have been deposited in the Electron Microscopy Data Bank under accession codes EMD-65597 [https://www.ebi.ac.uk/pdbe/entry/emdb/EMD-65597], EMD-65598 [https://www.ebi.ac.uk/pdbe/entry/emdb/EMD-65598], and EMD-65600 [https://www.ebi.ac.uk/pdbe/entry/emdb/EMD-65600]. The atomic models generated in this study have been deposited in the Protein Data Bank under accession codes 9W3D [https://doi.org/10.2210/pdb9W3D/pdb], 9W3E [https://doi.org/10.2210/pdb9W3E/pdb], and 9W3G [https://doi.org/10.2210/pdb9W3G/pdb]. The cryo-EM density map used in this study is available in the Electron Microscopy Data Bank under accession code EMD-8585 [https://www.ebi.ac.uk/pdbe/entry/emdb/EMD-8585]. The atomic models used in this study are available in the Protein Data Bank under accession codes 6C9Y [https://doi.org/10.2210/pdb6C9Y/pdb], 7MKP [https://doi.org/10.2210/pdb7MKP/pdb], and 7KHB [https://doi.org/10.2210/pdb7KHB/pdb]. Source data are provided with this paper.

## Acknowledgements

We thank Shenghai Chang at the Center of Cryo Electron Microscopy in Zhejiang University School of Medicine for help with cryo-EM data collection. We thank Cheng Ma from the Core Facilities, Zhejiang University School of Medicine for their technical support. We thank Zhijie Lin at Hangzhou Normal University for help with ITC experiments. This work was funded by National Key R&D Program of China (2023YFC2307100 to Y.F.), National Natural Science Foundation of China (32270030 and 32470019 to Y.F.).

## Author contributions

Yu Feng and Xiaoting Hua designed and supervised the study. Linggang Yuan, Qingyang Liu, Xiaojian Xiao, Liqiao Xu, Yang Guo, and Yue Yao performed the experiments. All authors contributed to the analysis of the data and the interpretation of the results. Linggang Yuan prepared all the figures. Yu Feng, Xiaoting Hua, and Youjun Feng wrote the manuscript with contributions from the other authors.

## Competing interests

The authors declare no competing interests.

## Notes

### Competing Interest Statement

The authors have declared no competing interest.

