## Supplementary Information for "AlphaFold 3-powered discovery of phage proteins that inhibit bacterial transcription"

Yuan et al.

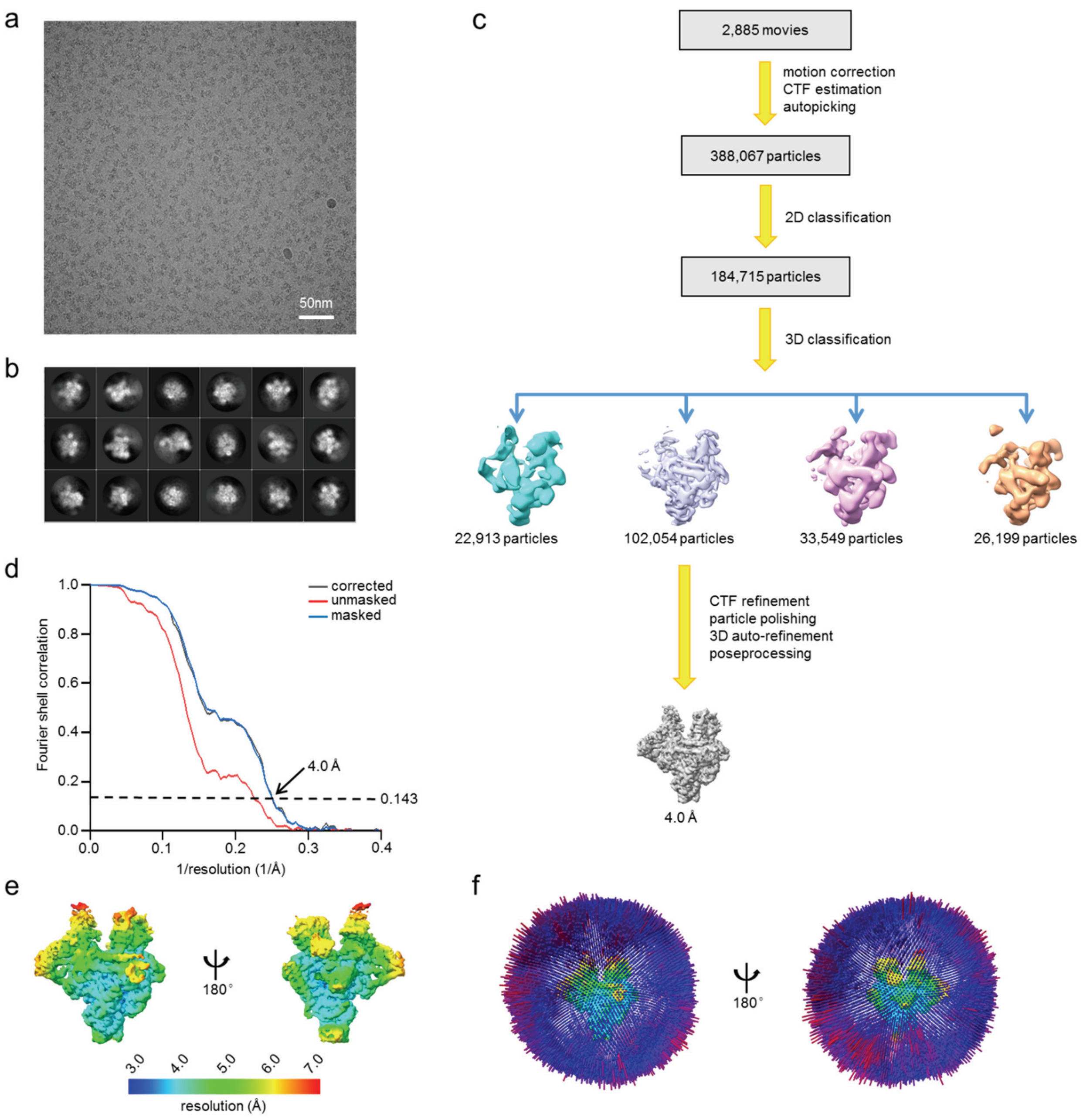


**Supplementary Fig. 1: Cryo-EM data processing and validation of RNAP-EP1.**

**a,** A cryo-EM micrograph of RNAP-EP1.

**b,** 2D averages of RNAP-EP1.

**c,** The cryo-EM data processing pipeline of RNAP-EP1.

**d,** The Fourier shell correlation (FSC) curves of RNAP-EP1. The gold-standard FSC is calculated by comparing the two independently determined half-maps from RELION. The dashed line represents the 0.143 cutoff.

**e,** The local resolution of RNAP-EP1.

**f,** The angular distribution of RNAP-EP1.

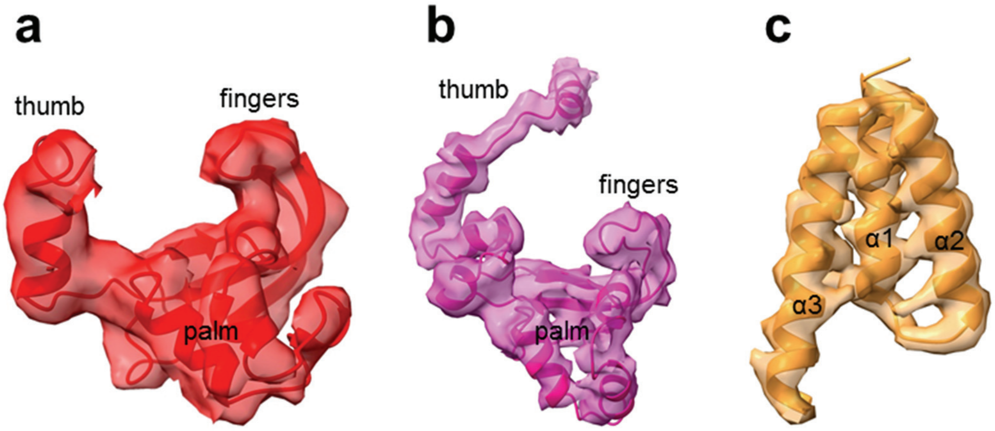


**Supplementary Fig. 2: Cryo-EM density maps of phage proteins.**

**a,** The map and model of EP1. The map without B-factor sharpening is shown as semi-transparent surfaces. The model of EP1 is shown as ribbons.

**b,** The map and model of PP1. The map without B-factor sharpening is shown as semi-transparent surfaces. The model of PP1 is shown as ribbons.

**c,** The map and model of VP1. The map without B-factor sharpening is shown as semi-transparent surfaces. The model of VP1 is shown as ribbons.

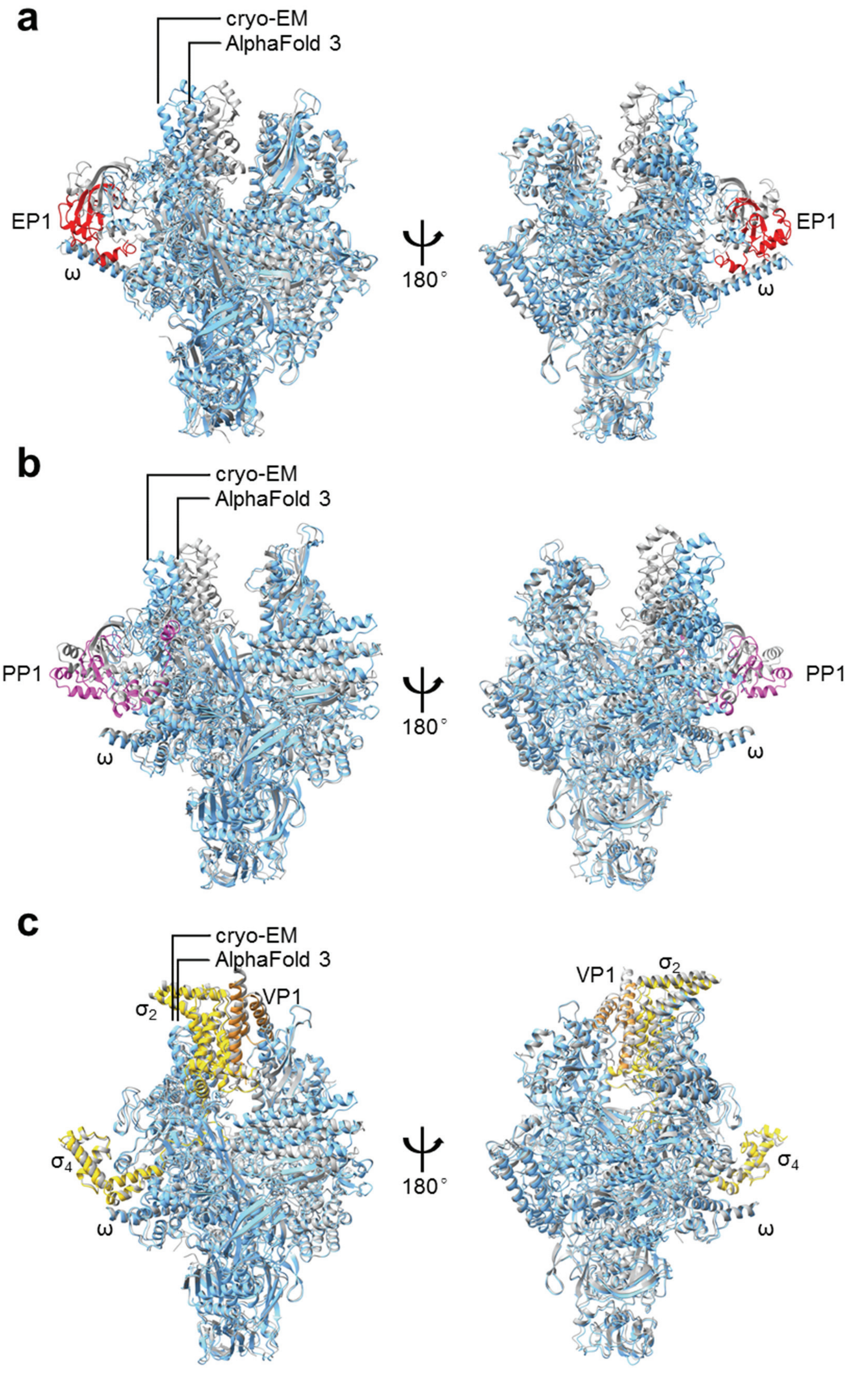


**Supplementary Fig. 3: Comparison of structures determined by cryo-EM and predicted by AlphaFold 3.**

**a,** Superimposition of RNAP-EP1 structures determined by cryo-EM (colored ribbons) and predicted by AlphaFold 3 (gray ribbons). Although AlphaFold 3 correctly predicted the folding of EP1 and its binding position, it failed to predict the rotation of the RNAP clamp and the EP1–ωCTH interaction.

**b,** Superimposition of RNAP-PP1 structures determined by cryo-EM (colored ribbons) and predicted by AlphaFold 3 (gray ribbons). Although AlphaFold 3 correctly predicted the folding of PP1 and its binding position, it failed to predict the rotation of the RNAP clamp.

**c,** Superimposition of RNAP-VP1 structures determined by cryo-EM (colored ribbons) and predicted by AlphaFold 3 (gray ribbons). The structure predicted by AlphaFold 3 closely matches the cryo-EM structure.

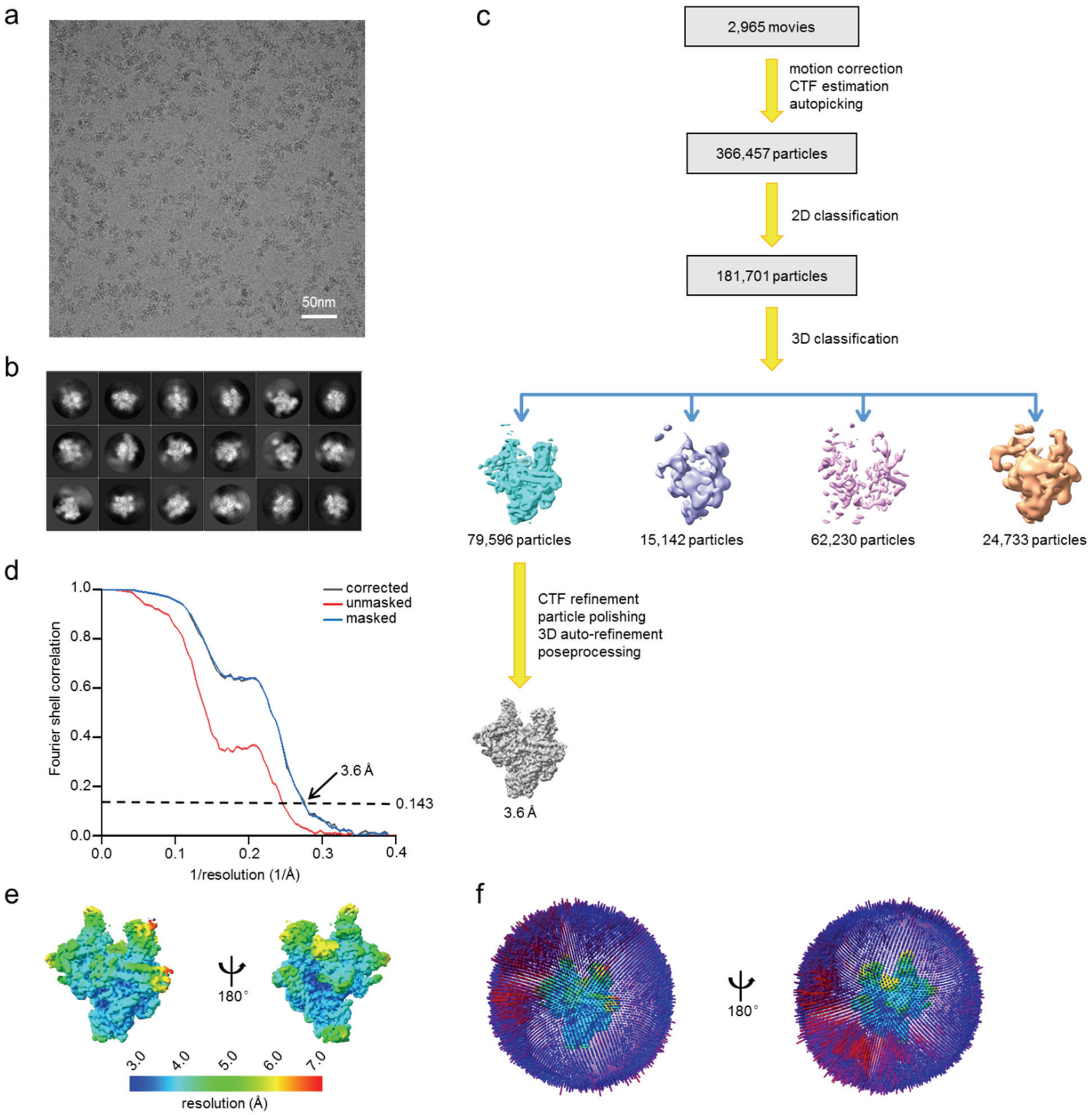


**Supplementary Fig. 4: Cryo-EM data processing and validation of RNAP-PP1.**

**a,** A cryo-EM micrograph of RNAP-PP1.

**b,** 2D averages of RNAP-PP1.

**c,** The cryo-EM data processing pipeline of RNAP-PP1.

**d,** The Fourier shell correlation (FSC) curves of RNAP-PP1. The gold-standard FSC is calculated by comparing the two independently determined half-maps from RELION. The dashed line represents the 0.143 cutoff.

**e,** The local resolution of RNAP-PP1.

**f,** The angular distribution of RNAP-PP1.

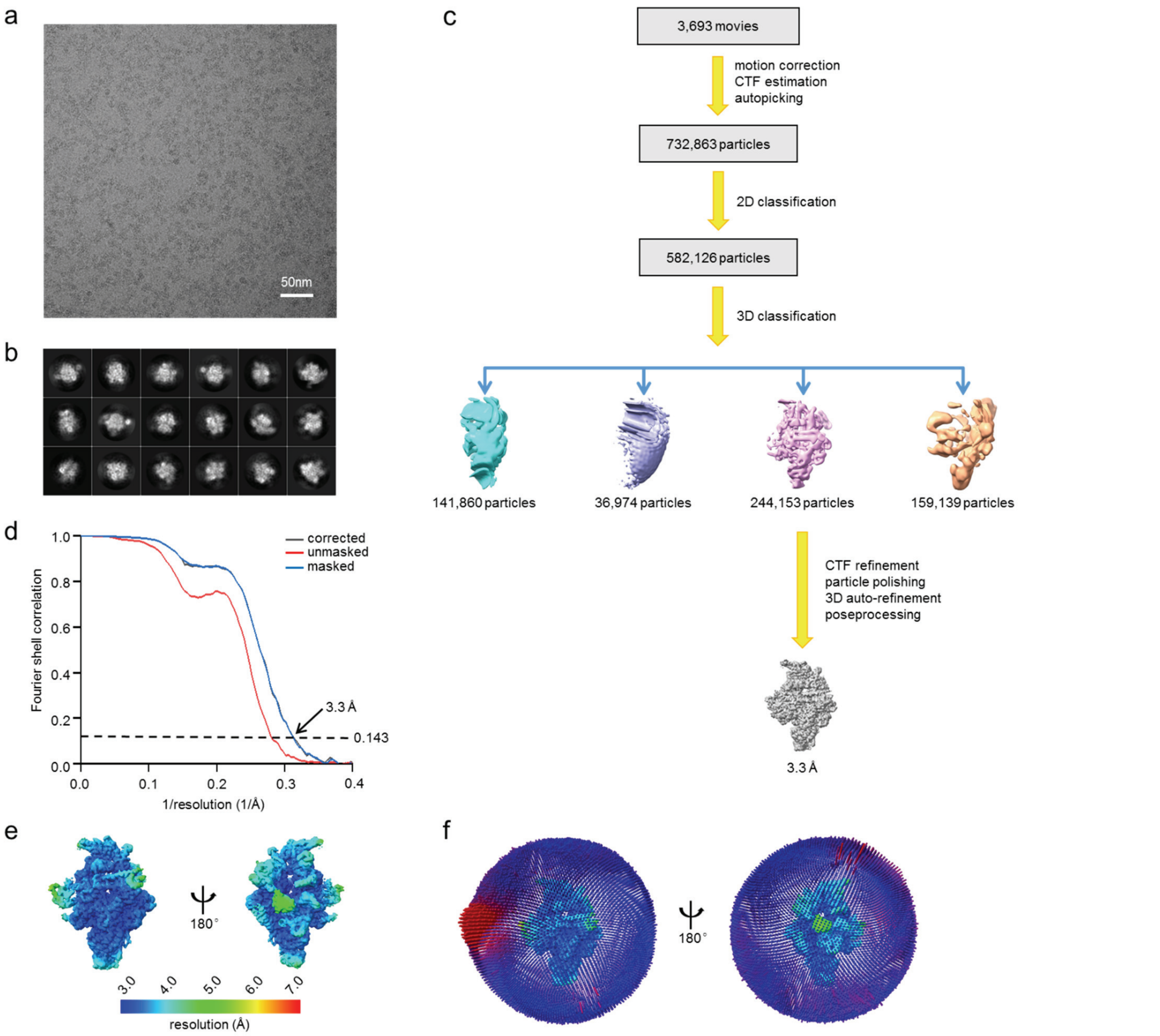


**Supplementary Fig. 5: Cryo-EM data processing and validation of RNAP-VP1.**

**a,** A cryo-EM micrograph of RNAP-VP1.

**b,** 2D averages of RNAP-VP1.

**c,** The cryo-EM data processing pipeline of RNAP-VP1.

**d,** The Fourier shell correlation (FSC) curves of RNAP-VP1. The gold-standard FSC is calculated by comparing the two independently determined half-maps from RELION. The dashed line represents the 0.143 cutoff.

**e,** The local resolution of RNAP-VP1.

**f,** The angular distribution of RNAP-VP1.

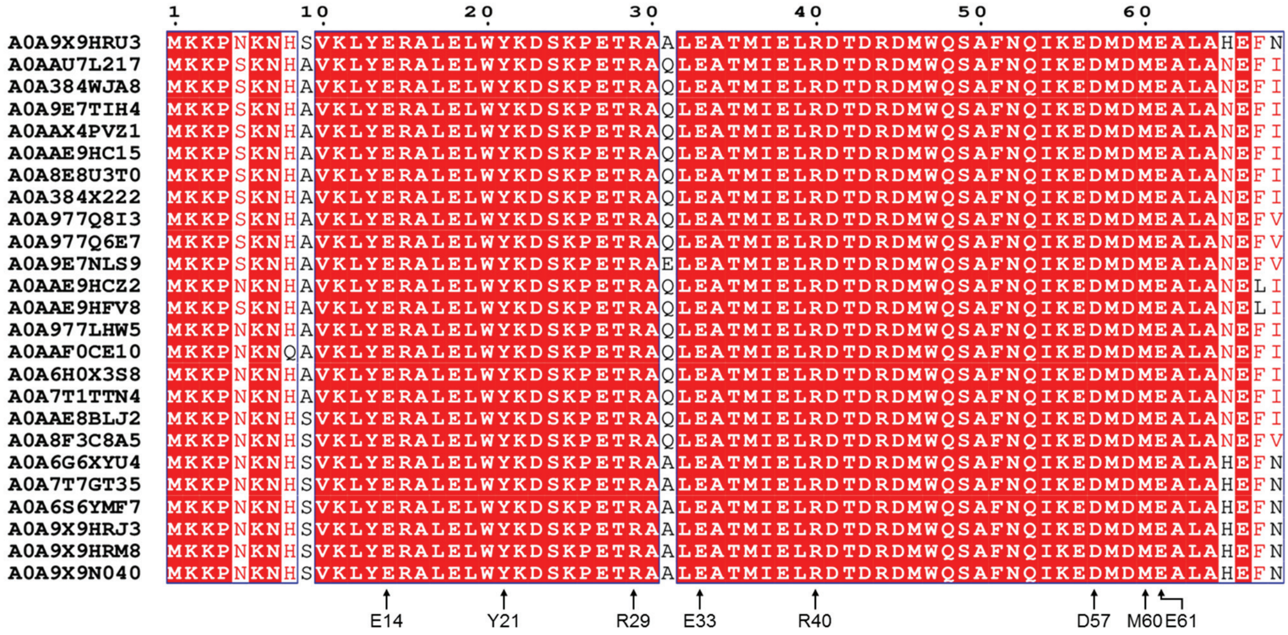


**Supplementary Fig. 6: Sequence alignment of VP1 homologs.**

Multiple protein sequence alignment was performed in Clustal Omega (https://www.ebi.ac.uk/jdispatcher/msa/clustalo), and the output was formatted with ESPript 3.0 (https://espript.ibcp.fr/ESPript/cgi-bin/ESPript.cgi). Contact residues discussed in this paper are highlighted by arrows. The UniProtKB IDs are given in front of the sequences. VP1 is the first protein in the alignment.

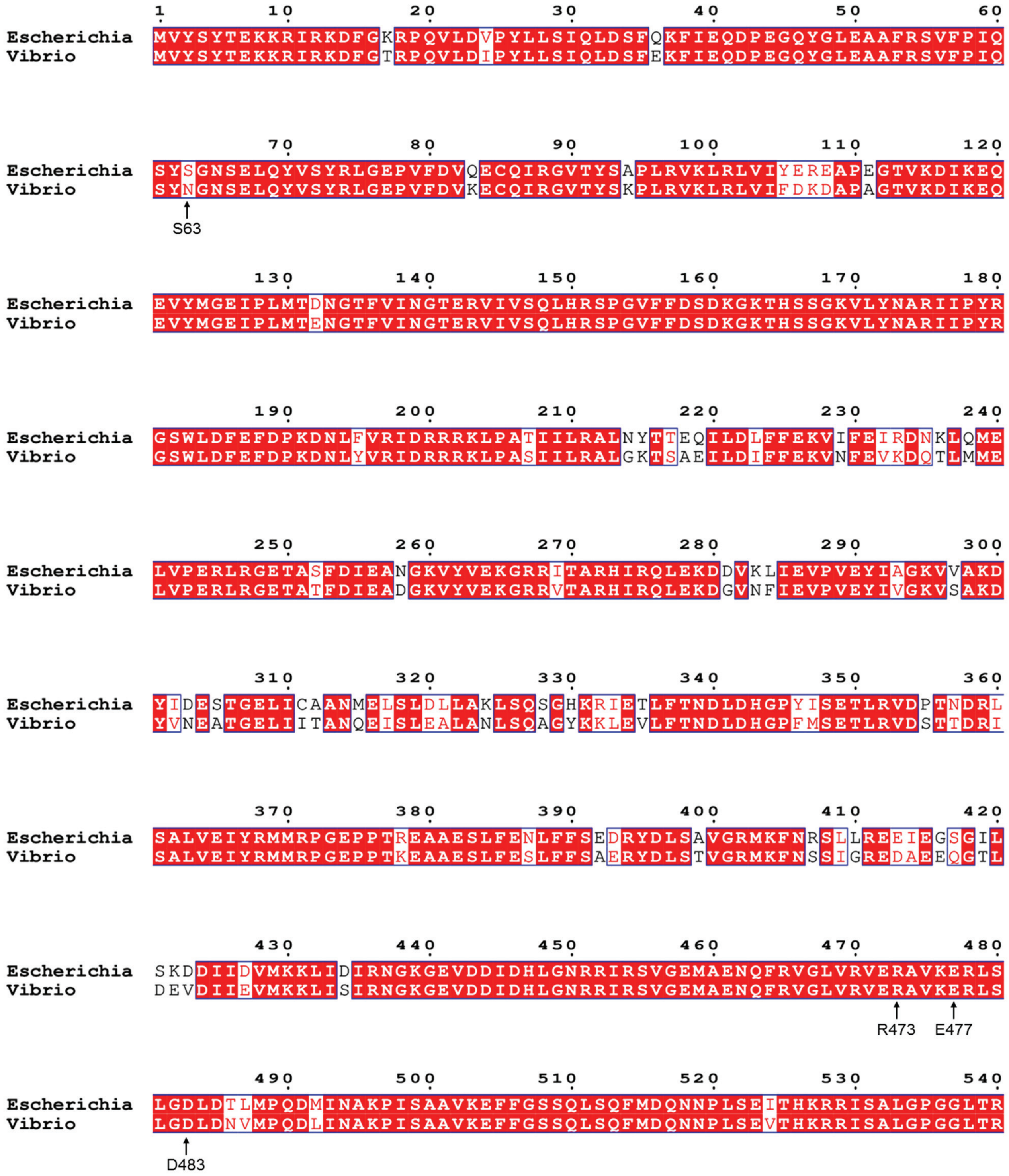


**Supplementary Fig. 7: Sequence alignment of *Escherichia* and *Vibrio* β subunits.**

Multiple protein sequence alignment was performed in Clustal Omega (https://www.ebi.ac.uk/jdispatcher/msa/clustalo), and the output was formatted with ESPript 3.0 (https://espript.ibcp.fr/ESPript/cgi-bin/ESPript.cgi). Contact residues discussed in this paper are highlighted by arrows. The UniProtKB IDs for *Escherichia* and *Vibrio* β subunits are P0A8V2 and Q9KV30, respectively. Only N-terminal 660 amino acids are shown due to space limitation.

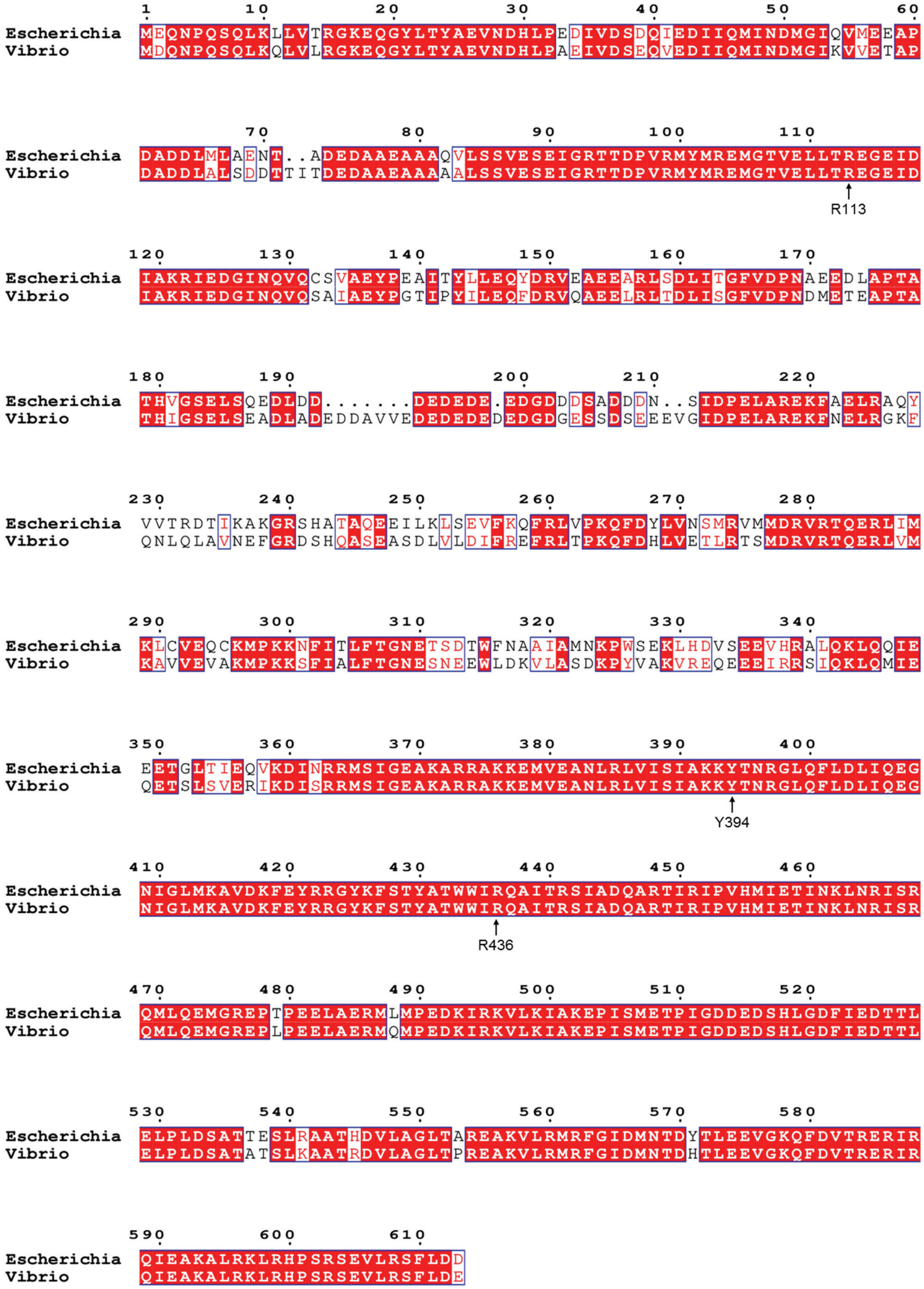


**Supplementary Fig. 8: Sequence alignment of *Escherichia* and *Vibrio* σ[70] factors.**

Multiple protein sequence alignment was performed in Clustal Omega (https://www.ebi.ac.uk/jdispatcher/msa/clustalo), and the output was formatted with ESPript 3.0 (https://espript.ibcp.fr/ESPript/cgi-bin/ESPript.cgi). Contact residues discussed in this paper are highlighted by arrows. The UniProtKB IDs for *Escherichia* and *Vibrio* $\sigma^{70}$ factors are P00579 and B7X6V9, respectively.

**Supplementary Table 4: Cryo-EM data collection and statistics**

| | RNAP-EP1 | RNAP-PP1 | RNAP-VP1 |
|---|---|---|---|
| **Data collection and processing** | | | |
| Microscope | Titan Krios | Titan Krios | Titan Krios |
| Voltage (kV) | 300 | 300 | 300 |
| Detector | Falcon 4 | Falcon 4 | Falcon 4 |
| Electron exposure (e/Å$^2$) | 50 | 50 | 50 |
| Defocus range (μm) | 1.0-2.0 | 1.0-2.0 | 1.0-2.0 |
| Data collection mode | Counting | Counting | Counting |
| Physical pixel size (Å/pixel) | 1.19 | 1.19 | 1.19 |
| Symmetry imposed | C1 | C1 | C1 |
| Initial particle images | 388,067 | 366,457 | 732,863 |
| Final particle images | 102,054 | 79,596 | 244,153 |
| Map resolution (Å)[a] | 4.0 | 3.6 | 3.3 |
| **Refinement** | | | |
| Root-mean-square deviation | | | |
| Bond lengths (Å) | 0.004 | 0.005 | 0.006 |
| Bond angles (°) | 0.887 | 0.846 | 0.833 |
| Molprobity statistics | | | |
| Clashscore | 6 | 6 | 4 |
| Rotamer outliers (%) | 0.08 | 0.40 | 0.29 |
| Cβ outliers (%) | 0.00 | 0.00 | 0.00 |
| Ramachandran plot | | | |
| Favored (%) | 97 | 97 | 98 |
| Outliers (%) | 0 | 0 | 0 |
| Map-to-model correlation coefficient | 0.86 | 0.86 | 0.80 |

[a]Gold-standard FSC 0.143 cutoff criteria
